# Pathogenic fungus exploits the lateral root regulators to induce pluripotency in maize shoots

**DOI:** 10.1101/2025.06.30.662278

**Authors:** Mamoona Khan, Nithya Nagarajan, Kathrin Schneewolf, Caroline Marcon, Danning Wang, Frank Hochholdinger, Peng Yu, Armin Djamei

## Abstract

Biotrophic plant pathogens secrete effector molecules to redirect and exploit endogenous signaling and developmental pathways in their favor. The biotrophic fungus *Ustilago maydis* causes galls on all aerial parts of maize. However, identification and characterization of the responsible gall-inducing effectors and corresponding plant signaling pathway(s) are largely elusive. Here we reveal molecular components acting downstream of a group of plant TOPLESS (TPL)- corepressor interacting fungal effectors that are directly involved in gall formation. We classify these effectors into two classes based on distinct phenotypic responses when individually expressed *in planta*. While class I responses are characterized by leaf chlorosis, transgenic plants expressing class II TPL-interacting protein (Tip) effectors show a derepression of the *AtARF7/AtARF19* branch of auxin signaling, resulting in pluripotent calli formation without the external addition of phytohormones. Subsequent comparative transcriptomics in maize also demonstrates a highly significant overlap of genes up-regulated during *U. maydis*-triggered leaf gall formation and during the developmental initiation of lateral root formation. This suggests that *U. maydis* can hijack the lateral root initiation pathway to induce galls in maize shoots. We demonstrate that underlying this is the transcriptional upregulation of downstream *LATERAL ORGAN BOUNDARIES DOMAIN (LBD)* transcription factors. Maize mutants in two LBD genes (*ra2, rtcs*) led to significantly reduced gall formation following *U. maydis* infection on homozygous mutants. This study reveals a set of dominantly acting cell proliferation-inducing fungal effectors and provides novel insight into plant developmental pathways that can be hijacked by biotrophic pathogens to induce gall formation.

## Introduction

Pathogen-induced plant galls are the morphological outcome of abnormal growth of plant tissue induced by the manipulative activities of the invading organism. These result from the increased proliferation (hyperplasia) and/or increased cell size of a group of cells (hypertrophy). Although more than a century of research, neither the physiological networks nor the exact mechanisms of gall induction and development have been fully elucidated (Dodueva et al., 2020). *Ustilago maydis*, a basidiomycete fungus, causes common smut disease in maize, infecting all aerial parts of the host, including the stem, leaves, and flowers. Early infection symptoms include local chlorosis and anthocyanin accumulation. A hallmark of *U. maydis* infection in maize is the formation of galls on above-ground organs, which act as sink tissues that, by the end of the fungal proliferation cycle, become filled with countless black diploid teliospores observable as smut symptoms (Brefort et al., 2009). *U. maydis*-induced gall formation results from intensive cell division and expansion in specific cell types (Matei et al., 2018). Notably, not all cells in the infected tissue proliferate, indicating that this process is cell-type or zone-specific. For instance, in maize flowers, *U. maydis* colonizes only immature, undifferentiated anther cells with meristematic activity, sustaining cell division beyond normal development (Gao et al., 2013; Lin et al., 2021). In leaf galls, differentiated bundle sheath cells resume cell divisions, and mesophyll cells enlarge (Matei et al., 2018). However, the molecular mechanisms involved in gall formation remain poorly understood.

The *U. maydis* genome encodes 467 predicted secreted proteins and several show distinct expression patterns during maize colonization (Lanver et al., 2018), however, only a few have been functionally characterized. Most known effectors are linked to immune suppression (Doehlemann et al., 2009; Navarrete et al., 2022; Navarrete et al., 2021; Saado et al., 2022), host metabolism, and phytohormone manipulation (Djamei et al., 2011; Ma et al., 2018; Rabe et al., 2016; Reineke et al., 2008; Tanaka et al., 2014) or fungal development in the host (Lin et al., 2023; Tanaka et al., 2020; Weiland et al., 2023). Yet little is known about effectors directly involved in gall formation. For example, See1 reactivates DNA synthesis in leaf galls but not tassel galls; its deletion inhibits hyperplastic cell division (Redkar et al., 2015). Sts2 promotes hyperplasia by transcriptional dysregulation, and its deletion reduces cell division in bundle sheath cells (Zuo et al., 2023). Another effector, ApB73, plays a cultivar-specific role in gall formation (Stirnberg and Djamei, 2016). The persistence of galls despite, the deletion of these effectors implies functional redundancy or cooperation among multiple pathways in this complex developmental process.

Auxin, a central regulator of plant growth, also plays a role in plant-pathogen interactions (Kunkel and Johnson, 2021; Nagarajan et al., 2023). During *U. maydis* infection, auxin levels increase, partly due to fungal INDOLE-3-ACETIC ACID (IAA) production, although this alone is not essential for gall formation (Reineke et al., 2008). Auxin-responsive genes are upregulated in infected tissues, and at least ten *U. maydis* effectors (Jsi1, Nkd1, Tip1–8) target TOPLESS (TPL) transcriptional co-repressors to modulate auxin signaling (Bindics et al., 2022; Darino et al., 2021; Huang et al., 2023; Khan et al., 2024; Navarrete et al., 2022). A pentuple deletion mutant of these (*Δtips1-5*) led to a significant reduction in gall sizes and numbers (Bindics et al., 2022), highlighting the crucial role of TPL proteins in the biotrophic stage of *U. maydis*. However, the specific downstream targets of TPL-controlled signaling remain to be explored. TPL corepressors in auxin signaling are recruited by AUXIN (Aux)/IAA proteins through conserved EAR domains (Szemenyei et al., 2008; Tiwari et al., 2004). Aux/IAAs do not bind DNA directly but interact with AUXIN RESPONSE FACTORs (ARFs) to suppress transcription. Since the ability of Aux/IAAs to modulate transcription is dependent on ARFs, the presence of different ARF complements in different cells can also affect auxin signaling specificity (Bargmann et al., 2013; Leyser, 2018; Rademacher et al., 2012).

Auxin also regulates pluripotency and organogenesis by regulating fate, division, and differentiation, enabling new organ formation throughout the plant lifecycle (Perianez-Rodriguez et al., 2014). Lateral root (LR) formation in plants is an example of postembryonic development. In *Arabidopsis thaliana* (*A. thaliana*) LRs originate exclusively from pericycle founder cells in response to local auxin maxima (Casimiro et al., 2001; Celenza et al., 1995), which results in the de-repression of AtARF7 and AtARF19, and expression of *LATERAL ORGAN BOUNDARY* (*LOB*) *DOMAIN* (*LBD*) transcription factors (TFs; Lee et al., 2009; Okushima et al., 2007; Wilmoth et al., 2005). Formation of callus, proliferating masses of pluripotent cells from various differentiated explants, is often the first step in *in vitro* plant generation and is also initiated by elevated auxin levels in the tissue culture medium (Lardon and Geelen, 2020). Intriguingly, callus formation involves an ectopic activation of the root primordia development program from pericycle or pericycle-like cells, even when derived from aerial organs, such as cotyledons and petals (Atta et al., 2009; Sugimoto et al., 2010; Xu et al., 2012). Auxin accumulation in the founder cells mediates the degradation of IAA14/SOLITARY ROOT (SLR), which releases AtARF7 and AtARF19 that in turn, upregulates the expression of *AtLBD16*, *AtLBD18*, and *AtLBD29* during callus formation (Lardon and Geelen, 2020). Ectopic LBD expression can induce callus without exogenous hormones, while their suppression inhibits auxin-induced callus formation (Fan et al., 2012; Okushima et al., 2005). The LBD family comprises 43 members in *A. thaliana* (Fan et al., 2012), and 49 in *Zea mays* (Zhang et al., 2020), however, only a few have been shown to regulate LR organogenesis and callus formation in *A. thaliana* (Feng et al., 2012; Goh et al., 2012; Lee et al., 2013; Pandey et al., 2018) and functional knowledge on LBDs in maize is limited. *Rootless concerning crown and seminal roots (rtcs)* and *rtcs-like (rtcl)* are two paralogous genes in maize and orthologues of *A. thaliana AtLBD29* (Berardini et al., 2015) that control seminal and post-embryonic shoot-borne root formation (Taramino et al., 2007; Xu et al., 2015). While *ramosa 2* (*ra2*) is an orthologue of *A. thaliana AtLBD25* and *AtLOB* genes (Zhang et al., 2020) and controls inflorescence branching (Bortiri et al., 2006). Although *ra2* is highest expressed in the roots of maize (Hoopes et al., 2019; Winter et al., 2007; Woodhouse et al., 2021), its direct role in LR formation in maize has not been explored.

In the present study, we reveal the role of a set of Tip effectors in *U. maydis-induced* cellular dedifferentiation, cell division, and gall formation. We show that a single Tip effector can induce cellular dedifferentiation leading to callus formation and cell division in transgenic *A. thaliana* plants. Furthermore, we provide genetic evidence that this process relies on the expression of *AtARF7* and *AtARF19* TFs. It is also demonstrated that *U. maydis* induces the expression of *Zmarf27*, an orthologue of *AtARF7* and *AtARF19,* and the LBDs which are dominant susceptibility factors in maize during biotrophic colonization. Transcriptomic overlap between gall formation and lateral root development supports the notion that *U. maydis* exploits parts of the lateral root formation pathway in maize for leaf-gall induction.

## Results

### Induction of Tip expression leads to strong morphological phenotypes *in planta*

Although *U. maydis* has at least ten TPL-interacting effectors (Jsi1, Nkd1, Tip1-8, (Bindics et al., 2022; Darino et al., 2021; Huang et al., 2023; Khan et al., 2024; Navarrete et al., 2022), it is not clear why so many are needed and which downstream pathways are activated. Since TPL co-repressors are central, conserved negative transcriptional regulators in all land plants, we generated transgenic *A. thaliana* lines expressing each of these Tip effectors to study their individual biological activity. Intriguingly, this resulted in a range of strong morphological phenotypes across two independent transgenic lines tested compared to an mCherry-expressing control (Figure 1, S1, S2 and S3). More specifically, the induction of *Tip1, Tip2, Tip8, Jsi1 and Nkd1* expression in *A. thaliana* led to chlorophyll loss in the cotyledons and leaves, and overall growth arrest already clearly visible at 4 days after transfer (Figure 1B, S1 and S2), while expression of *Tip3, Tip4, Tip5, Tip6,* and *Tip7* led to a strong induction of lateral root formation which was visible already two days after transfer to induction medium (data not shown, Figure 1C, S3). Noticeably, the TPL interacting effectors that lead to overall growth inhibition and complete chlorophyll loss in *A. thaliana*, previously showed a cell death phenotype upon transient overexpression in *Nicotiana benthamiana* leaves (Darino et al., 2021; Khan et al., 2024; Navarrete et al., 2022). Based on their cell death-inducing feature versus lateral root induction abilities, we categorized the ten known TPL interacting effectors into two classes, class I and II represented by Tip1 and Tip4 which exhibited the strongest phenotype of their respective class (Figure 1B &C). Strikingly, in the strongest case of class II effectors (Tip3, Tip4 and Tip6) the lateral roots did not grow in size, instead the whole root thickened due to the initiation of undifferentiated (callus-like) structures (Figure1C, 2A, B & D, E) that formed without the external addition of any phytohormones on the plate (Figure 1C and 2A, B, Figure S3 A, B & D). Surprisingly, this phenotype was restricted to the roots only; in the shoots, the growth either slowed down or the leaves turned pale. These plants continued to survive and the shoots could even undergo flowering (Figure 2C). To test if these undifferentiated (callus-like) structures are indeed pluripotent, we transferred them (without their shoots) to the shoot induction media used for *A. thaliana* regeneration. Impressively, this led to greening and shoot formation (Figure S4), supporting this conclusion. To summarize, ectopic expression of TPL interacting effectors in *A. thaliana* resulted in extreme morphological phenotypes, which could be grouped into two distinct classes. In this study, we focus on class II Tips whose expression leads to lateral root and callus induction in *A. thaliana* in-depth investigation.

**Figure 1.**
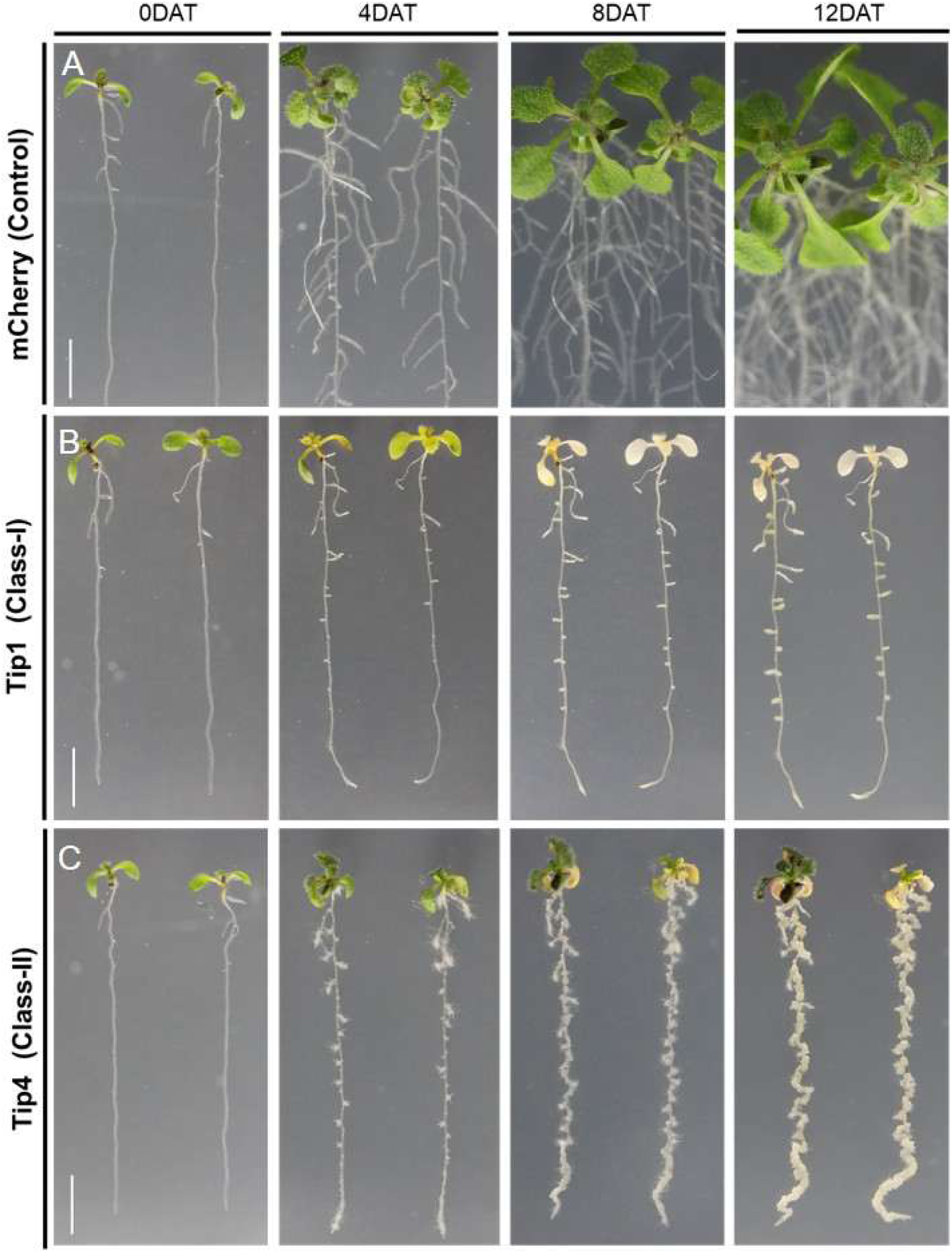
Expression of *TOPLESS* (TPL)-interacting protein (Tip) effectors induces strong morphological phenotypes in A. thaliana. Seven-day-old *A. thaliana* seedlings expressing (A) *pXVE: HA-mCherry*, (B) *pXVE: HA-mCherry-Tip1*, or (C) *pXVE: HA-mCherry-Tip4* were transferred to agar plates containing estradiol, and images were taken at 0, 4, 8, and 12 days after transfer (DAT). Scale bar = 1 cm.

**Figure 2:**
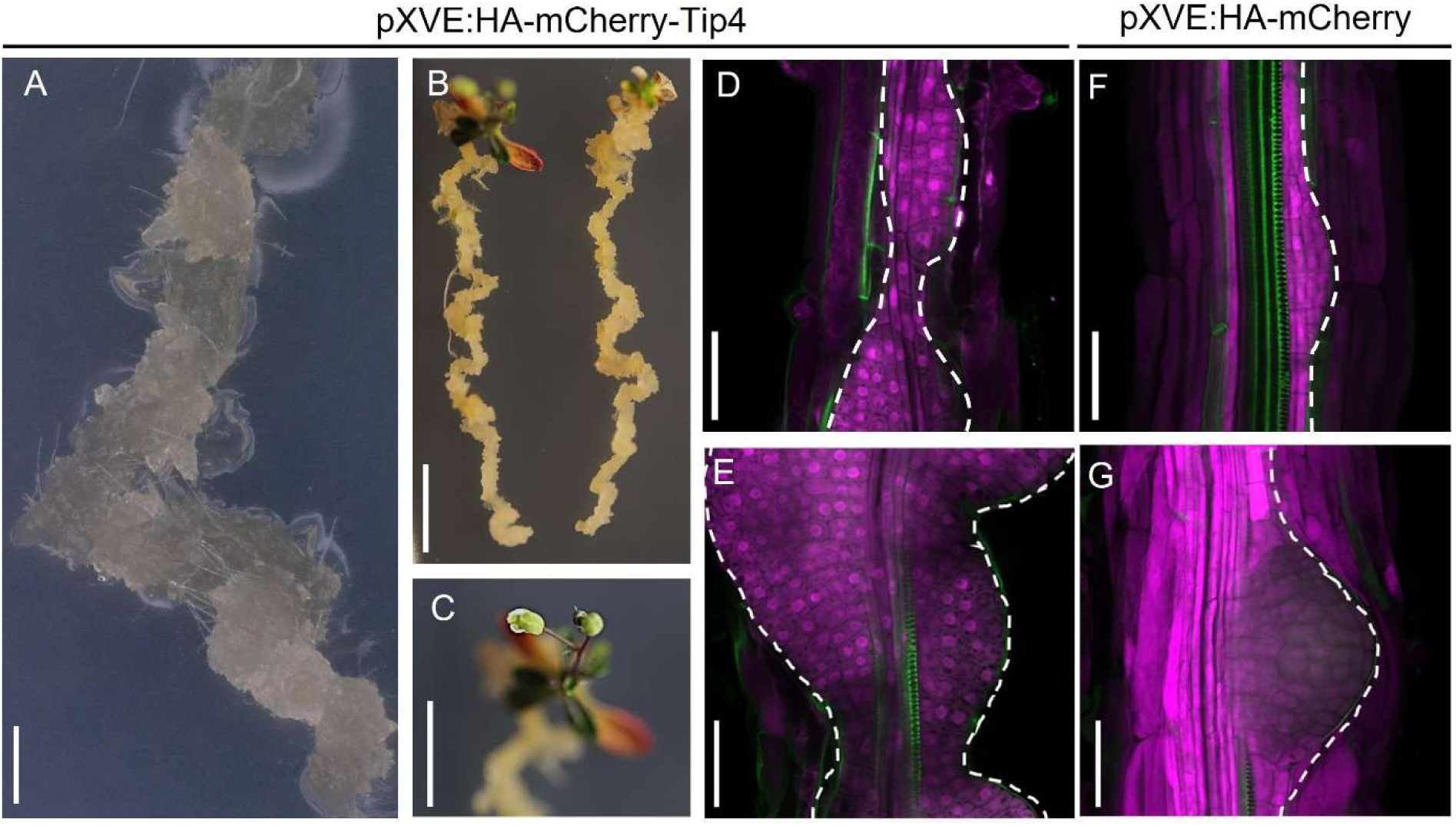
TOPLESS interacting protein (Tip) effector 4 induces callus-like structures along the primary root. **(A, B)** Digital microscopy images of plants expressing *pXVE:HA-mCherry-Tip4* at 12 days after transfer (DAT) (A) and 29 DAT (B) to estradiol induction medium, showing extensive callus formation along the primary root. (C) Flowering observed in a *pXVE:HA-mCherry-Tip4*-expressing plant at 29 DAT. Scale bars: A = 1 mm; B, C = 1 cm. (D–G) Confocal images of the root elongation zone in plants expressing *pXVE:HA-mCherry-Tip4* (D, E) and *pXVE:HA-mCherry* (F, G), showing distinct morphologies associated with callus formation (D, E) and lateral root development (F, G). Green fluorescence indicates auramine staining of the root vasculature. Scale bars: D–G = 50 µm.

### Overexpression of class II Tips leads to the formation of calli in *A. thaliana* roots through the TPL-AtARF7/AtARF19-AtLBD16 pathway

Expression of class II *U. maydis* Tip effectors (Tip3, Tip4, Tip5, Tip6, Tip7) led to the initiation of lateral roots and pluripotent callus in the *A. thaliana* roots. Similar phenotypes have been described previously for LBD transcription factor overexpression in *A. thaliana,* which acts downstream of AtARF7 and AtARF19 during lateral root formation and for callus formation (Fan et al., 2012; Liu et al., 2018). We therefore hypothesized that class II Tips, following their specific interaction with TPL class of co-repressors (Bindics et al., 2022; Navarrete et al., 2022), interfere with the binding of Aux/IAA corepressors and derepress the auxin signaling cascade associated with lateral root and callus formation in *A. thaliana.* To test this, we first crossed *pAtLBD16:GUS* (Bargmann et al., 2014) with the Tip4 expression line *pXVE:HA-mCherry-Tip4* (Tip4 representative for the class II phenotypes) and observed the GUS-reporter expression pattern. Microscopic observation revealed *pAtLBD16:GUS* expression was strongly induced in a Tip4-dependent manner in seedling roots as early as six hours (not shown) until at least four days after estradiol induction (Figure 3 A to L). This implicates that Tip-targeted TPLs are major negative regulators of *AtLBD16* in the root but there may be other factors in *Arabidopsis* shoots.

**Figure 3:**
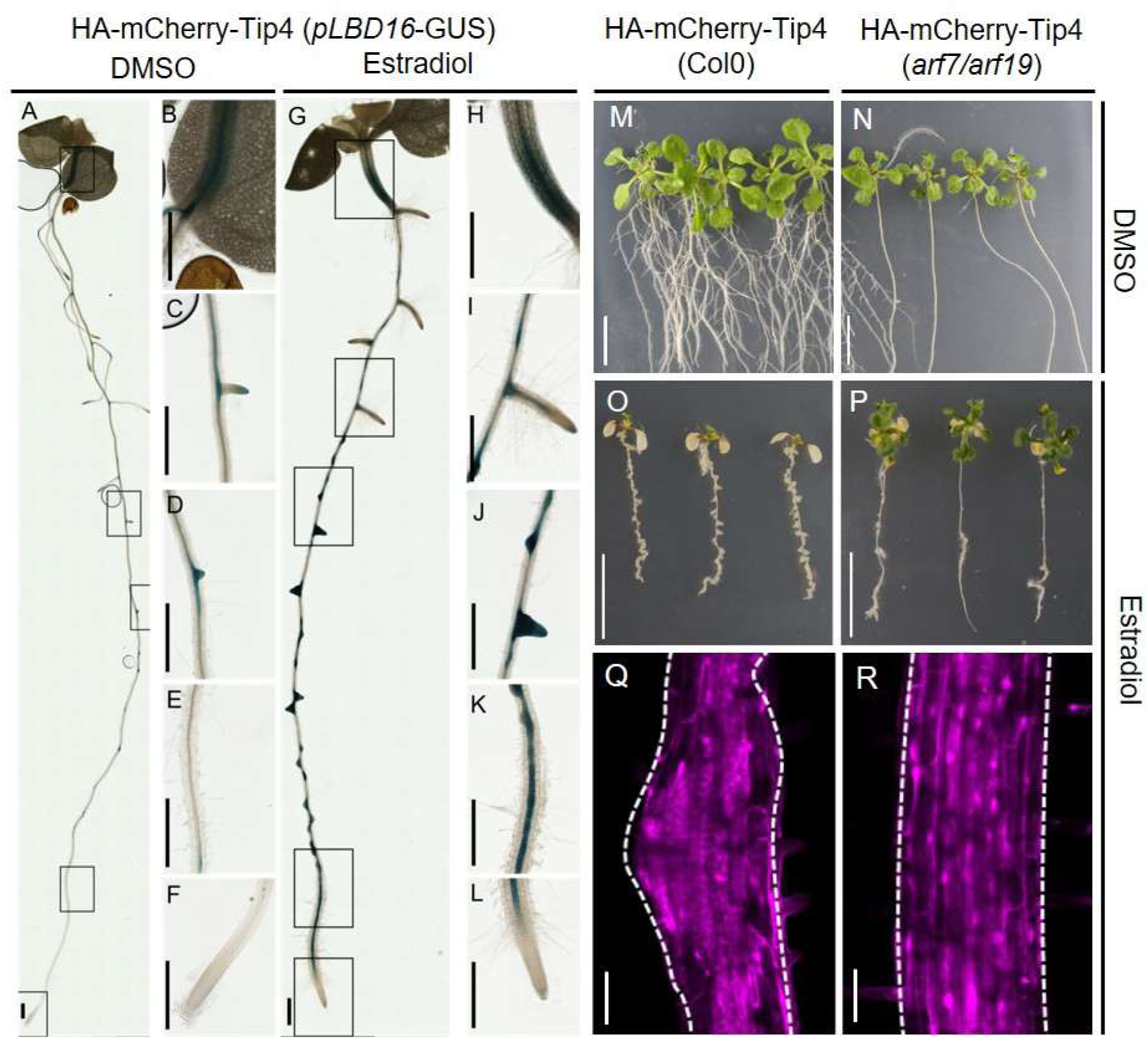
TOPLESS interacting protein (Tip)-4 effector–induced root callus formation requires AtARF7 and AtARF19–mediated AtLBD16 expression. **(A–L)** Digital microscopy images of seedlings expressing *pLBD16:GUS* in a *pXVE:HA-mCherry-Tip4* background. Seven-day-old seedlings grown on ½ MS agar were transferred to either DMSO-containing medium (A–F) or 10 µM estradiol (G–L) for 4 days before β-glucuronidase (GUS) staining. Scale bar = 500 µm. (M–P) Phenotypes of seedlings expressing *pXVE:HA-mCherry-Tip4* in wild-type Columbia (Col-0; M, O) or *arf7/arf19* mutant background (N, P) 10 days after transfer to plates containing DMSO (M, N) or 10 µM estradiol (O, P). Scale bar = 1 cm. (Q, R) Confocal microscopy of 11-day-old *A. thaliana* roots expressing *pXVE:HA-mCherry-Tip4* in Columbia (Q) or *arf7/arf19* background (R), showing effector protein expression 4 days after induction. Scale bar = 50 µm.

To provide further genetic evidence that morphological phenotypes of class II overexpressing effector lines are a result of the de-repression of a branch of the auxin signal cascade that controls lateral root and callus formation through *AtARF7* and *AtARF19*, we examined the overexpression phenotypes of Tip4 in an *arf7/arf19* T-DNA insertional double mutant background (Okushima et al., 2005) which does not produce lateral roots (Figure 3N). For this purpose, we crossed *pXVE:HA-mCherry-Tip4* (Figure 3M) expressing *A. thaliana* plants with the *arf7/arf19* double mutant (Figure 3N) and observed the phenotypes of homozygous *pXVE:HA-mCherry-Tip4 arf7/arf19* seedlings in the F2 generation with and without estradiol induction (Figure 3 O & P). As shown in Figure 3 M-R, whereas the Tip4-expression-mediated root length inhibition stayed unaffected, the Tip4-expression-mediated induced callus formation was largely abolished in the *arf7/arf19* background (Figure 3 O, &P). Furthermore, the effect on leaf chlorosis upon class II Tip expression is strongly reduced in the *arf7/arf19* mutant background (Figure 3 O, &P). This result suggests a direct role of *AtARF7* and *AtARF19* in both phenomena, the Tip effector mediated callus formation in *A. thaliana* via the lateral root pathway as well as the observed effects of chlorophyll loss in the leaf chloroplasts.

### *U. maydis* induces the expression of genes involved in lateral root formation during biotrophy in maize leaves

To estimate the role of the lateral root and callus formation pathway during *U. maydis* biotrophy on maize at the transcriptomic level, we compared previously published transcriptomic data of *U. maydis* infected maize leaf tissues at four days post-infection (dpi), the time point of gall formation (Lanver et al., 2018), with transcripts differentially regulated in phloem-pole pericycle cells, the cell-type which gives rise to lateral roots in maize (Figure 4A; Yu et al., 2016) (unpublished data). A comparison of these two data sets indicates several genes are commonly regulated during these two biological processes (Figure 4B & C). In total 28044 genes were expressed in sum of the two independent experiments (lateral root experiment and *U. maydis*-maize infection experiment). Among 20109 commonly expressed genes we found 3805 genes enriched in phloem-pole pericycle cells, whereas upon *U. maydis* infection 3809 maize genes are significantly (fold change > 2; FDR<0.05) induced at 4dpi. Between the two datasets of upregulated genes, 739 genes are upregulated in common, which is highly significant in a chi-square test for independence (p=1.739e-31; Figure 4D).

**Figure 4.**
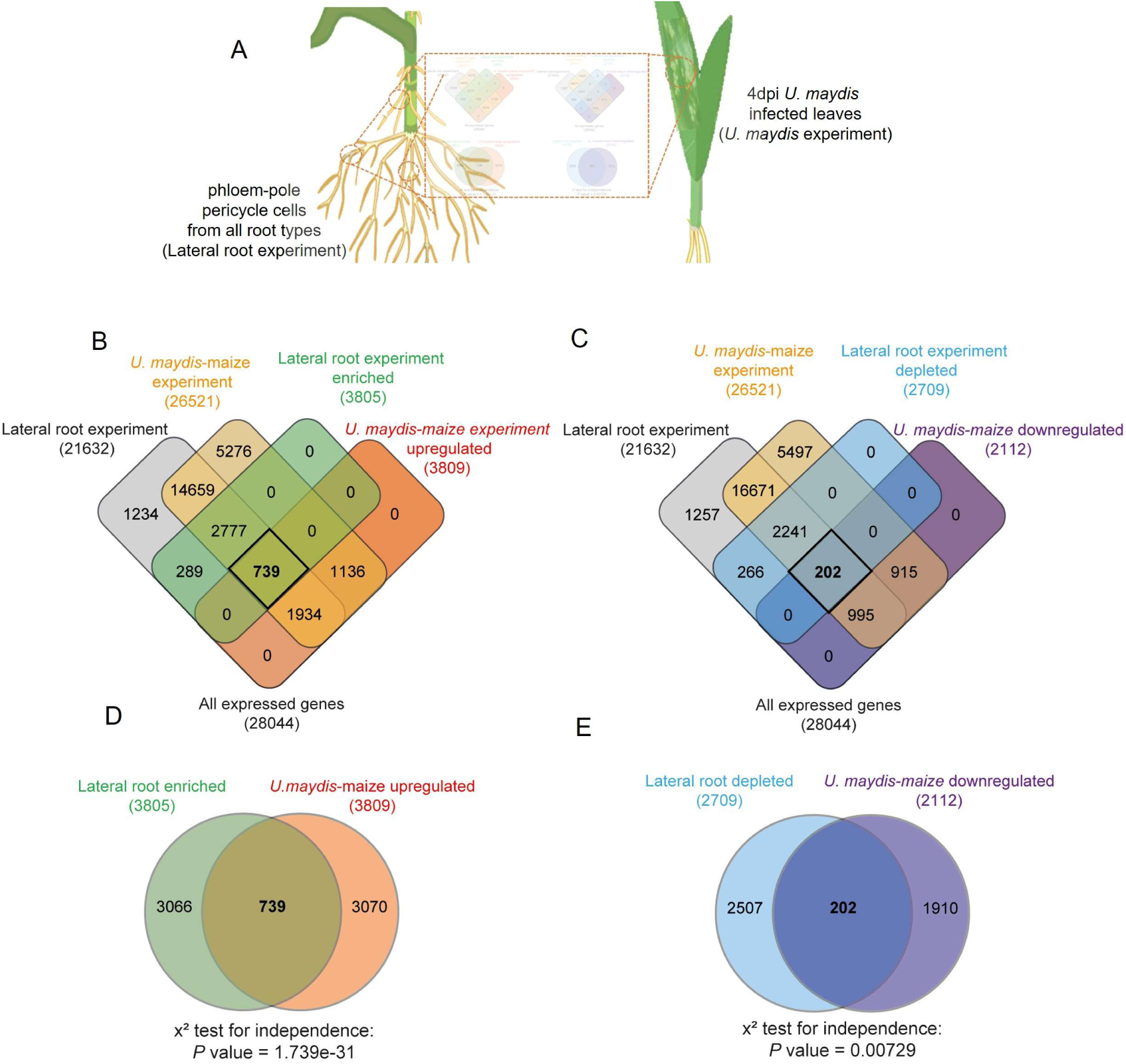
*Ustilago maydis* induces the expression of central regulators of the lateral root programming in maize shoots. (A) Schematic overview of the experimental design. Transcriptomic data from *U. maydis* infected maize leaf tissue at 4 days post-infection (4 dpi), the stage of gall formation, were compared with gene expression profiles from phloem-pole pericycle cells, the cell type responsible for LR initiation in maize, of 6-week-old plants. (B) 4-way Venn diagrams illustrating the overlap pattern of four different transcriptional datasets: expressed genes of lateral root (LR) experiment (21632), expressed genes of *U. maydis* maize infection experiment (26521), LR initiation enriched genes (3805) and *U. maydis* maize infection upregulated genes (3809) at 4dpi. (C) 4-way Venn diagrams illustrating the overlap pattern of four different transcriptional datasets: expressed genes of LR experiment (21632), expressed genes of *U. maydis* maize infection experiment (26521), LR initiation depleted genes (2709) and *U. maydis* maize infection downregulated genes (2112) at 4dpi. (D) Commonly upregulated genes between transcripts enriched in LR initiation cells and transcripts upregulated in 4dpi *U. maydis* infected maize. (E) Commonly downregulated genes between transcripts depleted in LR initiation cells and transcripts downregulated at 4dpi *U. maydis* maize infection. Statistical significance was assessed using the chi-square test for independence.

Subsequently, the 739 commonly induced genes between the datasets were functionally classified according to Gene Ontology (GO) terms using agriGOv2. In total, 69 GO terms belonging to different biological processes displayed significant overrepresentation (FDR < 0.05) (Supplemental Dataset 1). Of all GO terms, the most enriched are involved in processes of cell cycle and cell division, suggesting the potential linkage of lateral root initiation and *U. maydis* induced gall-formation in leaves. Also, the overlap of 202 commonly downregulated genes (Figure 4C) between the *U. maydis*-maize infection data and the lateral root initiation shows high significance in a chi-square test for independence (p=0.00729; Figure 4E). Analysis of the 202 genes (Figure 4C&E) from 2709 genes transcriptionally underrepresented in phloem-pole pericycle cells showed enrichment for 31 GOs (FDR < 0.05, Supplemental Dataset 1). In particular, these involved photosynthesis and chloroplast related pathways, consistent with sink-tissue formation and loss of photosynthetic activity at the place of gall-induction by *U. maydis*.

Taken together, comparison of the two independent transcriptomic data sets generated from very different tissues and with different biological questions shows a significant overrepresentation of commonly up and downregulated genes. This supports the notion that the shoot-infecting fungus *U. maydis* recruits the lateral root pathway during the induction of galls on maize shoots.

### *U. maydis* induces the expression of maize *Zmarf27*, the orthologue of *AtARF7* and *AtARF19,* in maize leaves

Next, we hypothesized that class II Tip effectors induce gall formation in maize leaves by employing common signaling components of the lateral root and callus formation pathway including *AtARF7/AtARF19* and LBDs. Therefore, we searched for maize orthologues of these pathway genes and identified one gene (*Zmarf27*) as the orthologue of both *AtARF7* and *AtARF19* and two genes (*Zmlbd24* and *Zmlbd1*) as orthologues of *AtLBD16* (Figure 5A; Berardini et al., 2015). We then examined their expression levels in *U. maydis* infected maize leaves compared to mock control by quantitative real-time PCR (qRT-PCR). We also included the *AtLBD29*, an orthologue *rtcs*, and *ra2,* a closest homologue of class-IB LBD (Figure 5A) genes of *A. thaliana. Ra2* was previously found to be downregulated in pentuple-deletion-mutant (*Δtips1-5*) infected plants (Khan et al., 2024). Moreover, ra2-like binding sites were enriched in differentially-expressed genes from *Δtip6* mutant-infected maize leaves (Huang et al., 2023). Strikingly, the expression levels of *Zmarf27, ra2,* and *Zmlbd1* were significantly and specifically induced in the leaves infected with *U. maydis* (Figure 5B). At the same time, *rtcs* was consistently upregulated but not significantly different between *U. maydis* infected leaves compared to mock treatment in our assays. The transcript abundance of *Zmlbd24* was below detection limits in both *U. maydis-infected* and mock leaves (data not shown).

**Figure 5.**
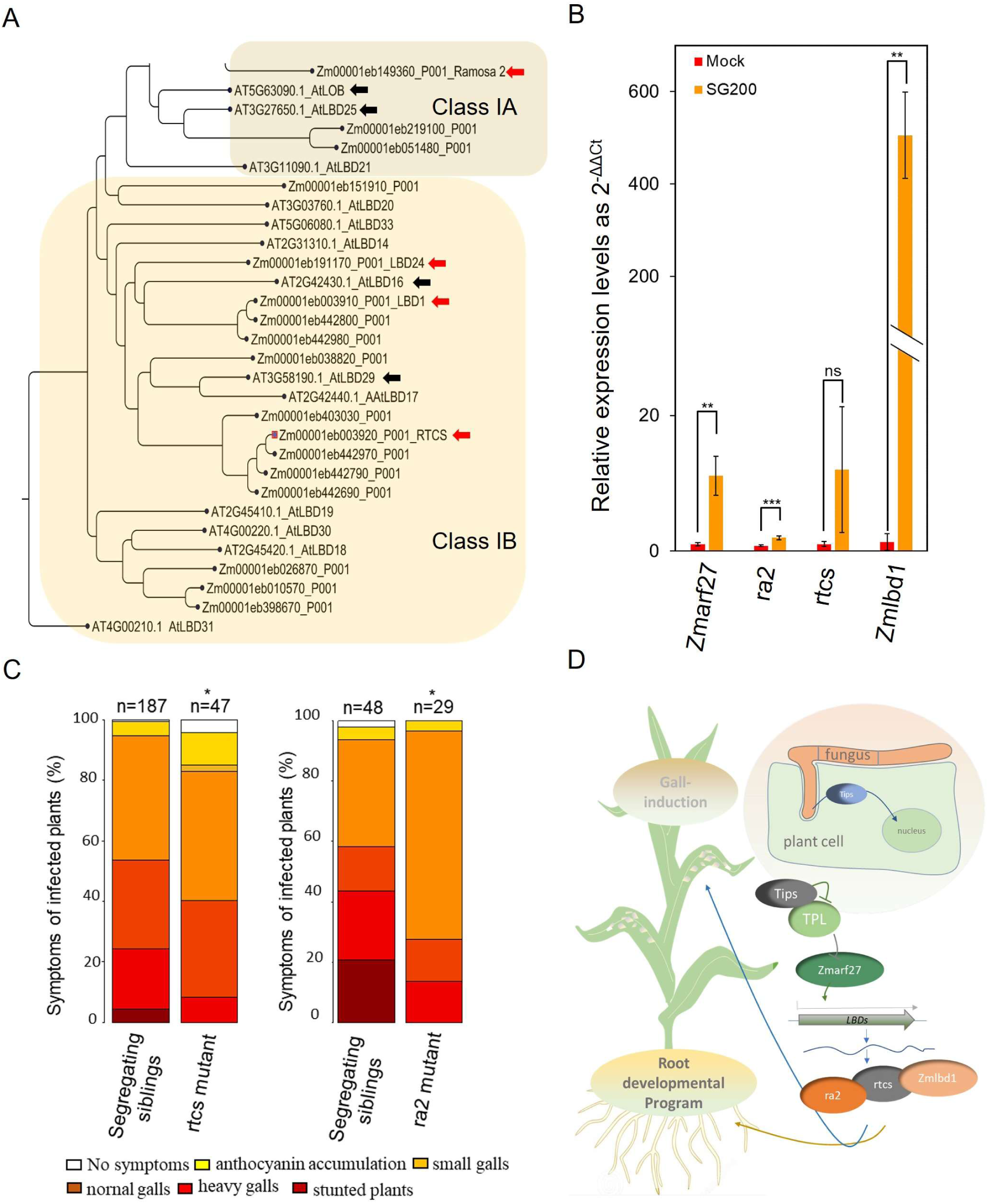
*Zmarf27* and certain ZmLBDs (*ra2, rtcs, Zmlbd1*) are required for *U. maydis-induced* gall formation. **(A)** The phylogeny of LOB domains (LBD) proteins of class-IA and class IB of *Arabidopsis thaliana* and *Zea mays*. The phylogenetic tree was reconstructed using aligned amino acid sequences of LOB domain proteins as an unrooted tree using CLC-Genomics software 20.04 and a bootstrap value of 1000. Red arrows point to maize genes studied in the fig. 5B-D and black arrows point to their *A. thaliana* orthologues. (B) The expression levels of *Zmarf27*, *ra2*, *rtcs,* and *Zmlbd1* were tested by quantitative real-time PCR analyses at four days post-infection of *Ustilago maydis* SG200 solopathogenic strain or mock (water) infected maize seedlings. The 2^−ΔΔCt^ values were calculated and plotted as relative expression levels compared with mock-infected plants as negative control. Error bars indicate standard deviation. Significant differences between mock control and SG200 infected leaves were analysed by Student’s t-test analysis (* = p< 0.05, ** = p < 0.01, *** = p < 0.005, ns, not significant). Data represent an average of four biological replicates. (C) Nine-day-old recessive homozygous maize mutants in *rtcs* oe *ra2* genes were infected with *Ustilago maydis* solopathogenic strain SG200 and compared to segregating population controls. Significant differences were analysed by Fischer’s exact test (*= p < 0.05). (D) Working model for class II TPL interacting protein effectors induced cellular reprogramming in maize shoot: Class II TPL-interacting protein (Tip3, Tip4, Tip5, Tip6 & Tip7) effectors interact with maize TPL proteins during *U. maydis* biotrophy in maize leading to de-repression of auxin response factor 27 (Zmarf27), which in turn can bind to the promoters of certain *LATERAL ORGAN BOUNDARIES DOMAIN* (LBD) transcription factors promoting the transcriptional reprogramming towards cellular dedifferentiation and gall formation in the shoot while their normal developmental role is in the root morphogenesis.

To elucidate the role of this pathway during *U. maydis* biotrophy, we next looked for the maize mutants of the respective LOB-domain transcription factors and identified previously published mutants in *rtcs* and *ra2* genes. We tested the ability of the recessive *rtcs* and *ra2* loss-of-function mutants to form galls upon *U. maydis* infection. For this purpose, 9-day-old maize seedlings of segregating *rtcs-1* and *ra2-R* alleles were infected with the solopathogenic *U. maydis* strain SG200, and symptom scoring was performed twelve days post-infection. The results of this experiment show a slight but significant reduction in virulence in homozygous recessive mutants compared to the segregating population (Figure 5 C&D). Taken together, *U. maydis* induces the expression of *Zmarf27*, and certain LBDs during its biotrophy in maize leaves and single *rtcs* and *ra2* mutants show virulence defects suggesting a role of this branch of auxin signaling during gall formation. Moreover, the weaker virulence defects of recessive single mutants of LBD genes indicate their redundant roles in this process.

## Discussion

More than a century of *U. maydis* research has been in-part driven also by the curiosity of how this fascinating biotrophic fungus can induce prominent galls on all aerial parts of its host plant maize (Küster and 1911). However, the underlying mechanisms remained elusive. Since the *U. maydis* genome has been published (Kamper et al., 2006) many effector genes have been placed into the context of gall formation based on studies of their deletion strains, which were impaired in the pathogenicity and hence altered gall formation upon infection (Brefort et al., 2014; Djamei et al., 2023; Khan and Djamei, 2024; Lanver et al., 2017; Redkar et al., 2015). Nevertheless, it was unknown how many factors would be needed to induce galls *in planta* in the absence of fungal infection. Here we provide significant insights through *in planta* by Tip effector overexpression, where major phenotypic changes were observed and could be classified in two distinct classed. For class II Tip effectors (*Tip3, Tip4, Tip5, Tip6,* and *Tip7*) we provide genetic evidence that they de-repress *AtARF7* and *AtARF19* transcription factors, leading to activation of *LBD* genes involved in lateral root and callus formation. In contrast to the previously reported effectors, the class II Tip effectors (Tip3, Tip4, Tip5, Tip6, and Tip7) act in a dominant fashion and independent of the presence of *U. maydis* infection, where overexpression of a single effector is sufficient to induce an endogenously encoded cellular dedifferentiation and proliferation program. This leads to the observed pluripotent callus-tissue formation where lateral roots are normally formed. We further show an overlap in transcriptional programming between *U. maydis* induced gall formation in maize leaves and lateral root emergence. These results are completely novel in *U. maydis-Zea mays* pathosystem and in line with recent findings in *A. thaliana* that demonstrate auxin-induced callus formation occurs from the pericycle (or pericycle-like cells) within multiple organs through a root development pathway, during which the ectopic activation of root meristematic genes is required for subsequent regeneration programs (Che et al., 2007; Sugimoto et al., 2010).

The discrepancy that *U. maydis* causes galls occur solely on the aerial organs of its host plant maize, while class II Tips overexpressed in *A. thaliana* solely induce calli in the roots, requires some consideration. One explanation could be that during *U. maydis* infection, the complex background manipulation by a whole effectome creates the metabolic and transcriptional precondition. This would include de-repression of *AtLBD16* homologs as demonstrated in Figure 3. Additionally, the ability to create these preconditions seems to be restricted, as *U. maydis* cannot induce cell division (hyperplastic leaf gall-tissue) in the whole leaf tissue. but only within specific cell types, i.e., bundle sheet cells (Matei et al., 2018). Nevertheless, class II Tips show clear effects also in the shoot of transgenic plants, i.e., chlorosis. This situation also occurs in *U. maydis* galls, which are sink tissues that show a loss of chloroplasts and turn pale yellow or red due to anthocyanin accumulation.

It has been demonstrated previously in *A. thaliana* that the formation of LRs, as well as the development of callus, both require several common regulatory components of the LR developmental pathways, i.e. *AtARF7, AtARF19, AtLBD16, AtLBD18, AtLBD29, and AtLBD33* (Fan et al., 2012; Perianez-Rodriguez et al., 2014; Sugimoto et al., 2010). The formation of callus upon *Tip4* overexpression was almost abolished in *arf7/arf19* double mutant background, placing *AtARF7* and *AtARF19* downstream of Tip4 and demonstrating their importance in Tip4-induced callus formation in *A. thaliana.* The fact that there were still a few callus-like structures appearing upon *Tip4* overexpression in the *arf7/arf19* mutant background could be due to the involvement of other ARFs e.g. *AtARF5* in this process (Vangheluwe and Beeckman, 2021). Leaves of *A. thaliana* plants expressing class II Tips turn chlorotic and this phenomenon seems also to be dependent on *AtARF7* and *AtARF19* as the leaf phenotypes of Tip4 are widely rescued in the *arf7/arf19* mutant plants (Figure 3O, P). We also observed this in western blots targeting class II Tips overexpression after already five days induction. This correlates with the yellowing of the leaf and the rubisco band in the ponceau-staining of western blots are very faint (data not shown), indicating that not only chlorophyll but chloroplast function in total is negatively regulated by AtARF7 and AtARF19 activity. Consistent with this, GO-enrichment analysis of the commonly downregulated genes between *U. maydis* infected maize leaves and lateral root initiation repressed genes revealed an overrepresentation of photosynthesis and chloroplast related components.

Gall formation is a widely occurring phenomenon caused by various pathogens like gall wasps, bacteria like Phytoplasmas, *Pantoea agglomerans*, *Pseudomonas savastanoi*, *Xanthomonas citri*, *Rhodococcus fascians*, the root-knot and cyst nematodes, and certain rust fungi and smuts (Harris and Pitzschke, 2020). Gall formation might have several advantages for the colonization and proliferation in the host including immune suppression (Navarrete et al., 2022) and efficient nutrient acquirement (Horst et al., 2010; Sosso et al., 2019). Furthermore, considering the biotrophic lifestyle of these pathogens, there is possibly also a reduction of interference with essential plant functionalities during massive proliferation in a separated tissue irrelevant for plant survival. We provide here evidence that *U. maydis* employs the post-embryonic organogenesis pathway to induce galls. Unlike galls and giant cells formed by root-knot nematodes, where direct overexpression of *AtLBD16* is induced for callus formation, the class II Tip effectors of *U. maydis* induce callus formation upstream of AtARF7 and AtARF19 by distinct suppression of TPL functions. This highlights an example of convergent evolution of the pathway comprising TPL as a negative regulator of AtARF7 and AtARF19.

## Supporting information

supplementary data Khan et al.

## Acknowledgments

The research leading to these results received funding from the European Research Council under the European Union’s Seventh Framework Programme ERC-2013-STG, Grant Agreement: 335691, the Austrian Science Fund (FWF): [P27818-B22, I 3033-B22], the Austrian Academy of Sciences (OEAW), and the Deutsche Forschungsgemeinschaft (DFG, German Research Foundation) under Germany’s Excellence Strategy EXC-2070-390732324 and DFG grants (DJ_64/5-1, DJ_64/7-1). We thank Prof. Dr. Gabriel Schaaf for critical reading of the manuscript and useful suggestions. We thank Dr. Evan John for proofreading and valuable feedback and fruitful discussions on the manuscript. We thank Dr. Ruben Betz for the fruitful discussions. We thank the technical staff and greenhouse facility of INRES / University of Bonn for plant care.

## Author contributions

Conceptualization: MK, AD. Methodology: MK, AD. Investigation: MK, NN, KS. Transcriptomic analysis DW, PY. Project administration: MK. Resources: MK, NN, CM, FH, AD. Original Draft: MK, AD. Funding acquisition: AD, Supervision AD, MK

## Materials and methods

### Molecular cloning

Cloning was performed using either Greengate (Lampropoulos et al., 2013) or Golden gate (Katzen, 2007) cloning systems as described previously (Khan et al., 2024). Mach1 competent cells (Thermo Fisher Scientific, Waltham, MS, USA) were used for all DNA manipulations and were grown in a dYT liquid medium or on YT agar plates with the required antibiotic supplements. *pXVE-HA-mCherry-(effector lacking signal peptide)* was the construct used for generating transgenic effector lines.

### Plant material and growth conditions

*A. thaliana* Columbia was a wild type used for generating all transgenic lines. Maize EGB was used for *U. maydis* infection assays for qRT-PCR analysis. *pAtLBD16:GUS* reporter line and T-DNA insertional mutant *arf7,arf19* lines were obtained from NASC under the Nasc ID N68141 and N24629 respectively and *ra2-R* maize mutant seeds were received from maize genetic and s cooperation stock center (USA). *A. thaliana* for dipping were grown in controlled short-day conditions (8h light/16h dark at 21°C ± 2 °C) while maize was grown in distinct conditions (14 h light/10 h dark, 28 °C/20 °C). Floral dipping was used to generate two transgenic lines for each plasmid construct with similar protein levels, selected on glufosinate-ammonium. For phenotyping, the plants were grown on solidified half-strength MS agar media, supplemented with 1% (w/v) sucrose plates in growth cabinets (Panasonic environmental test chamber, Type: MLR-352H-PE) at 21°C ± 2°C and on long days of 16 h light// 8 h dark cycles with 80 µmol/m^2^/s intensity. For estradiol treatments, 7 day-old plate-grown seedlings were transferred either to DMSO or 10µM β-estradiol containing ½ MS plates, and pictures were taken 5, 10, 15, and 20 days after transfer to the plates or otherwise stated. Experiments were repeated at least 3 times.

### Accession numbers

Zmarf27 (Zm00001eb373970), Zmlbd1 (Zm00001eb003910), Zmlbd24 (Zm00001eb191160), rtcs (Zm00001eb003920), ra2 (Zm00001eb123060), Tip1 (UMAG_11415), Tip2 (UMAG_02535), Tip3 (UMAG_02537), Tip4 (UMAG_02538) and Tip5 (UMAG_11417), Tip6 (UMAG_11060), Tip7 (UMAG_05300), Tip7 (UMAG_05308), Jsi1 (UMAG_001236), Nkd1 (UMAG_02299), AtARF7 (AT5G20730), AtARF19 (At1g19220), AtLBD16 (At2g42430), AtLBD25 (AT3G27650), AtLOB (At5g63090).

### Maize infection assays

Maize variety Early Golden Bantam, Old Seeds, Madison, WI, USA was used for all infections until otherwise stated. *U. maydis* progenitor strain SG200 was used to infect seven-day-old maize seedlings for qRT-PCRs and nine-days-old maize seedlings for mutant analyses as described in detail previously (Redkar and Doehlemann, 2016). For qRT-PCRs leaf samples were collected approximately 1 cm below the hole of the syringe in infected leaves 4-days-post-infection whereas symptom scoring was performed at 12 dpi according to (Kamper et al., 2006). Symptom scores were assessed using the Fisher exact test in R, as described previously (Stirnberg and Djamei, 2016).

### Confocal microscopy

Confocal microscopy was performed with a Leica SP8 confocal microscope. mCherry was excited at 561 nm and emission was collected between 578-648 nm. Images were processed using the LAS-X software from Leica.

### qRT-PCR and comparative transcriptome analyses

mRNA was extracted from the ground powder of leaves using a New England Biolabs GmbH (Frankfurt am Main, Germany) RNA extraction kit and cDNA was synthesized using Thermo Scientific RevertAid First Strand cDNA Synthesis Kit. qPCRs were performed with GoTaq qPCR mix (NEB, cat. no. A6001) according to the manufacturer’s instructions. Relative amounts of amplicons were calculated according to the 2^−ΔΔCt^ method (Livak and Schmittgen, 2001). The results are the average of four biological replicates. *Zmcdk* (Zm00001eb350890) was used as housekeeping gene (Lin et al., 2014).

### Biostatistical analysis of lateral root dataset and *U. maydis* dataset

To understand the biological pathway during lateral root initiation and callus formation in maize, a previously published transcriptomic dataset of *U. maydis* infected maize leaf tissues (Lanver et al., 2018) was downloaded and only the time point four days post-infection was selected as it is the time where galls are formed. The lateral root initiation dataset was derived from a previous study, which used cell-type specific RNA sequencing of lateral root mutant versus wild type plants via laser capture microdissection (Yu et al., 2016); (Unpublished dataset).FCDK Genes preferentially expressed in phloem-pole pericycle cells were defined as ‘lateral root enriched’, while genes expressed in xylem-pole pericycle cells were defined as ‘lateral root depleted’. DESeq2 was used to test for gene expression changes. For both datasets, only genes with at least five normalized read counts for at least one timepoint/cell type from three replicates were considered as expressed. A differential expression threshold of log_2_ fold change > 1 and Benjamini-Hochberg-adjusted *P* value < 0.05 was used in both datasets. The overlapped expressed and differentially expressed genes between datasets were visualized by Venn diagram (https://bioinformatics.psb.ugent.be). A Chi-square (χ^2^) test in R (version 1.4.1717) was used to determine if the two transcriptome datasets share a significant correlation. The overlapping genes between datasets were functionally characterized by gene ontology (GO) annotation and enrichment analysis via AgriGO (version 2, http://systemsbiology.cpolar.cn/agriGOv2/).

