## supplementary data Khan et al. for "Pathogenic fungus exploits the lateral root regulators to induce pluripotency in maize shoots"

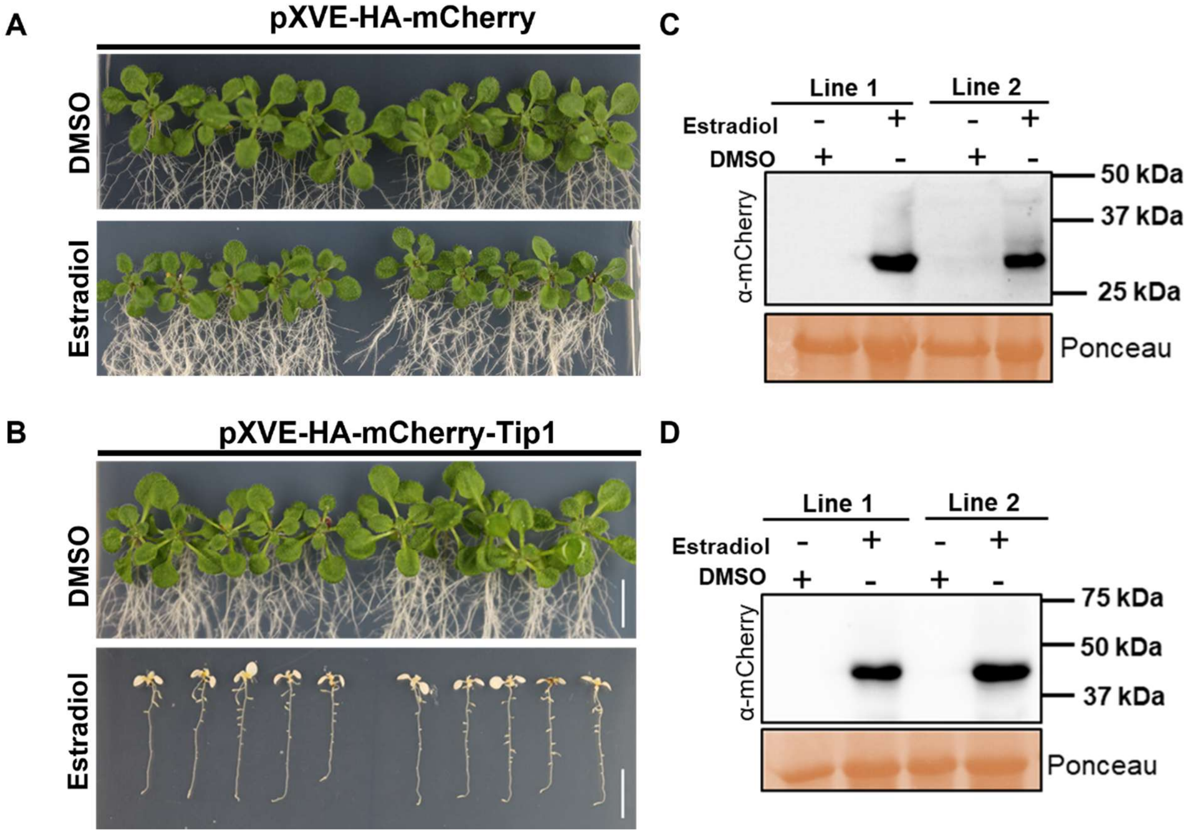


**Supplementary Figure 1:** ***Arabidopsis thaliana*** **plants expressing Topless interacting protein effector 1 (Tip 1) show chlorophyll loss, and inhibition of overall growth phenotypes.** Seven-day-old ½ MS agar-grown seedlings expressing (A) *pXVE:HA-mCherry* (B) *pXVE: HA-mCherry-Tip1* were moved to either DMSO or 10µM estradiol containing ½ MS agar plates, and images were taken at 10 days after the transfer. In each panel, right and left are two independent lines, scale bar = 1cm (C, D) Western blot analysis of total protein extracts of 8-day-old plants treated with either DMSO or 20µM estradiol for 2 hours. Membranes were incubated with α-mCherry antibody. Ponceau staining shows loading control.

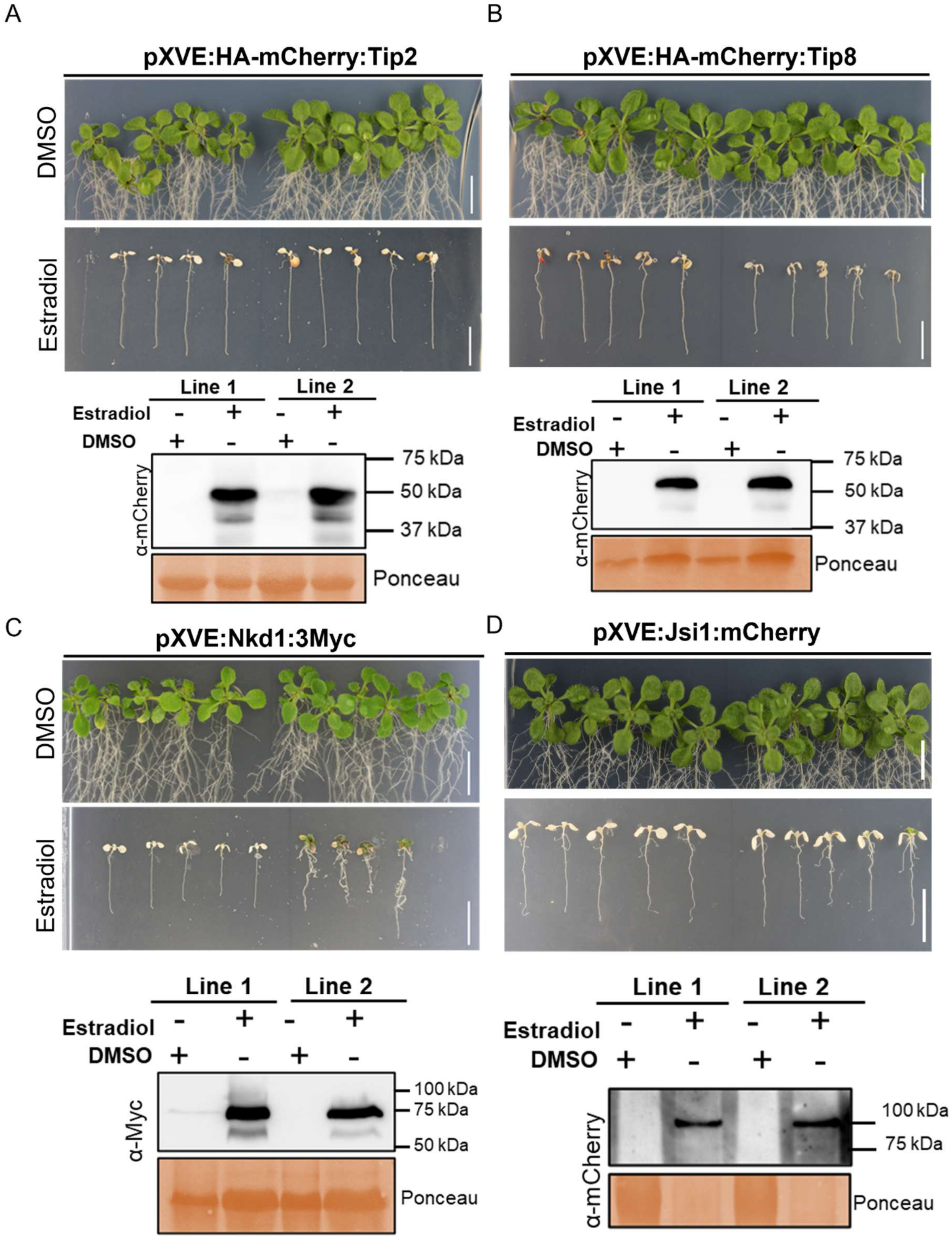


**Supplementary Figure2:** ***Arabidopsis thaliana*** **plants expressing Topless interacting protein (Tip) effectors of class I showing chlorophyll loss, and inhibition of overall growth**

**phenotypes.** Seven-day-old ½ MS agar-grown seedlings expressing (A) *pXVE:HA- mCherry-Tip2* (B) *pXVE:HA-mCherry-Tip2* (C) *pXVE:Nkd1-3myc* and (D) *pXVE:Jsi-mCherry* were moved to either DMSO or 10μM estradiol containing ½ MS agar plates and images were taken at 10 days after the transfer. In each panel, right and left are two independent lines, scale bar = 1cm (C) Western blot analysis of total protein extracts of 8-day-old plants treated with DMSO or 20μM estradiol for 2 hours. Membranes were incubated with α-mCherry antibody. Ponceau staining shows loading control.

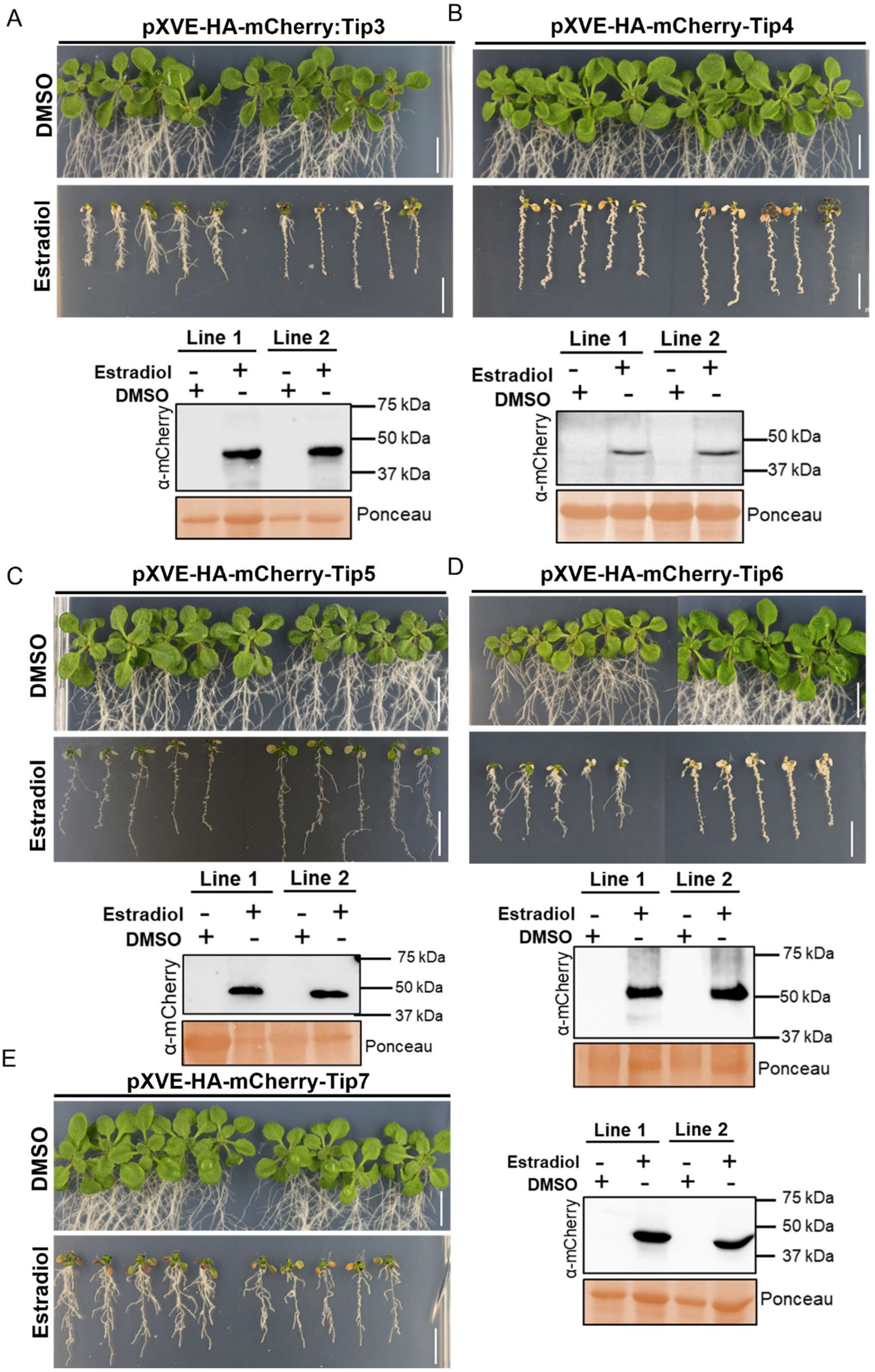
A
pXVE-HA-mCherry:Tip3
DMSO
Estradiol
Line 1
Line 2
Estradiol - + - +
DMSO + - + -
α-mCherry
75 kDa
50 kDa
37 kDa
Ponceau
B
pXVE-HA-mCherry-Tip4
DMSO
Estradiol
Line 1
Line 2
Estradiol - + - +
DMSO + - + -
α-mCherry
50 kDa
37 kDa
Ponceau
C
pXVE-HA-mCherry-Tip5
DMSO
Estradiol
Line 1
Line 2
Estradiol - + - +
DMSO + - + -
α-mCherry
75 kDa
50 kDa
37 kDa
Ponceau
D
pXVE-HA-mCherry-Tip6
DMSO
Estradiol
Line 1
Line 2
Estradiol - + - +
DMSO + - + -
α-mCherry
75 kDa
50 kDa
37 kDa
Ponceau
E
pXVE-HA-mCherry-Tip7
DMSO
Estradiol
Line 1
Line 2
Estradiol - + - +
DMSO + - + -
α-mCherry
75 kDa
50 kDa
37 kDa
Ponceau

**Supplementary fFigure 3:** ***Arabidopsis thaliana*** **plants expressing Topless interacting protein (Tip) effectors of class II showing phenotypes of increased lateral/ callus-like structure and inhibition of root length.** Seven-day-old ½ MS agar-grown seedlings expressing (A) *pXVE:HA-mCherry-Tip3* (B) *pXVE:HA-mCherry-Tip4* (C) *pXVE:HA-mCherry-Tip6* (D) *pXVE:HA-mCherry-Tip7* and (E) *pXVE:Nkd1-3xMyc* were moved to either DMSO or 10µM estradiol containing ½ MS agar plates, and images were taken at 10 days after the transfer. In each panel right and left are two independent lines, scale bar = 1cm (C) Western blot analysis of total protein extracts of 8-day-old plants treated with either DMSO or 20µM estradiol for 2 hours. Membranes were incubated with either α-mCherry antibody (for Tip3, Tip4, Tip6, Tip7) or with α-Myc antibody (Nkd1). Ponceau staining shows loading control.

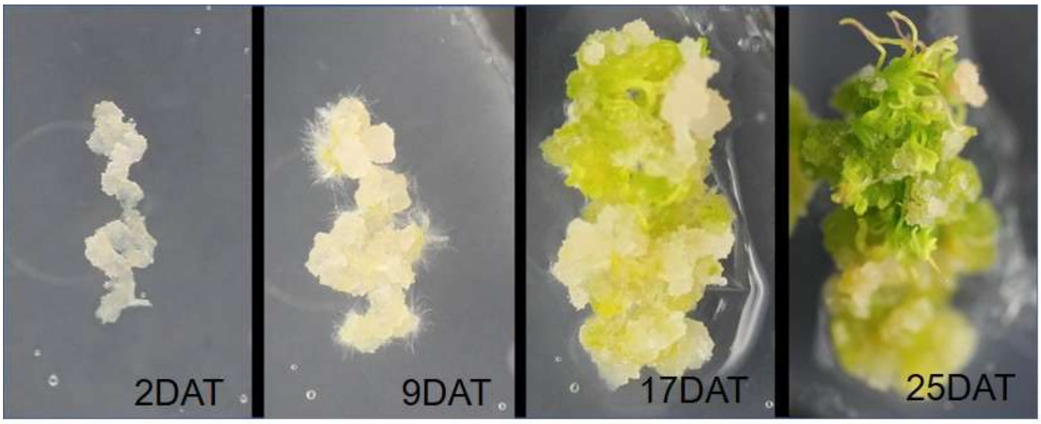


**Supplementary Figure 4**: The root explants of *pXVE:HA-mCherry-Tip4* pre-incubated with 10µM estradiol for the induction of Tip4 expression were transferred to shoot-inducing medium (SIM) to induce de novo shoot regeneration. The SIM plates were incubated under long day conditions and images were taken 2 days after the transfer (DAT), 9 DAT, 17 DAT, and 25 DAT to SIM.

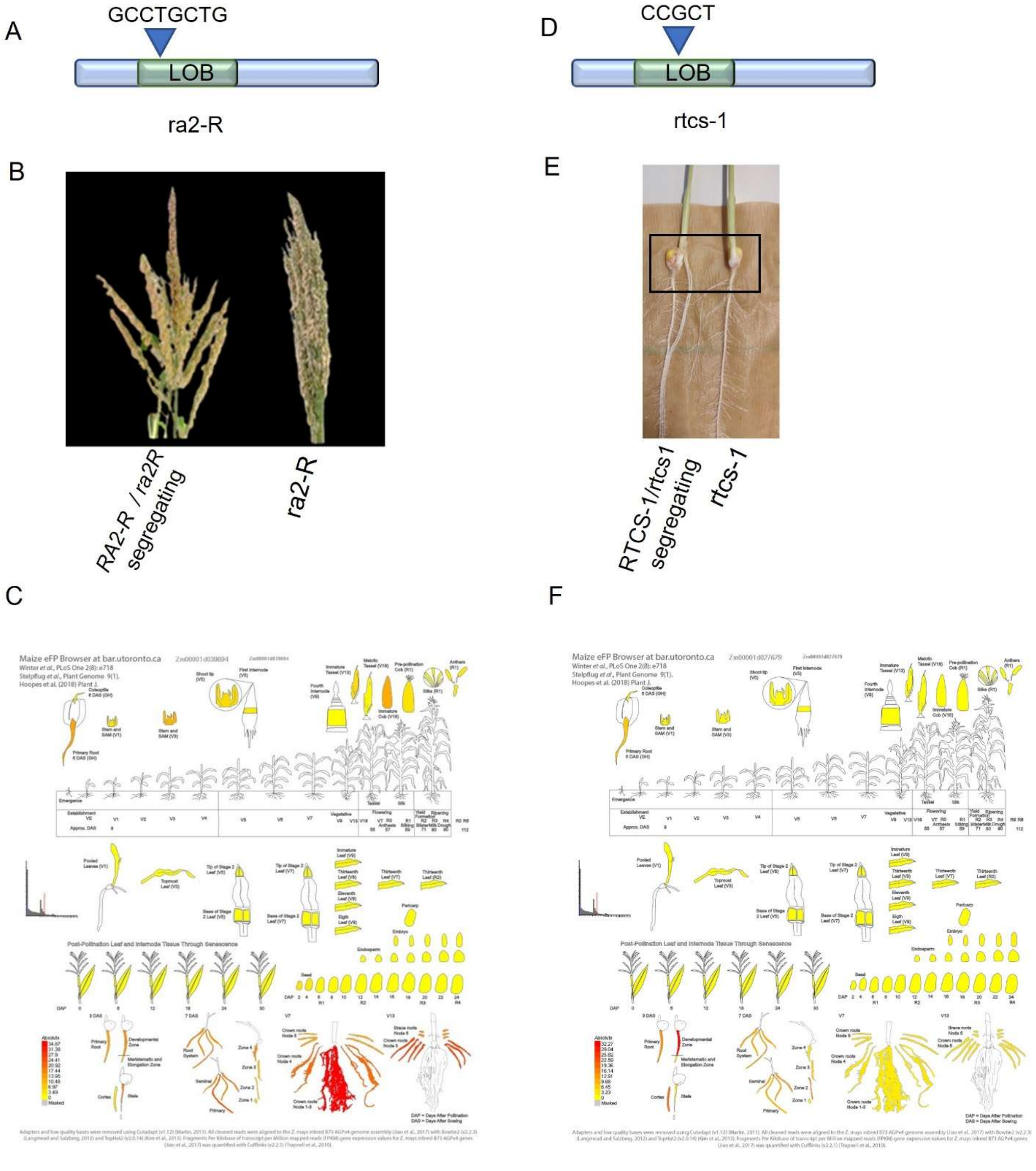


**Supplementary Figure 5**: **Characterization of *ra2-R* and *rtcs-1* recessive mutants.** (A) The *ra2-R* allele has an eight-base-pair (bp) insertion in the LATERAL ORGAN BOUNDARIES (LOB)- DOMAIN of *Ramosa2* (*RA2*) that introduces a stop codon within the LOB domain. (B)

The tassel phenotype of *ra2-R* mutant compared to the segregating control. (C) Tissue-specific expression of *RA2* (Zm00001d039694) according to (Hoopes *et al.*, 2019; Woodhouse *et al.*, 2021). (D) The *rtcs-1* allele has a five-base-pair insertion in the LATERAL ORGAN BOUNDARIES (LOB)- DOMAIN of *RTCS* that introduces a stop codon 227 bp downstream of the putative ATG start codon. (E) The phenotype of 8-day-old *rtcs-1* mutant; it does not form shoot-borne roots compared to the segregating control. (F) Tissue-specific expression of *RTCS* (Zm00001d027679) according to (Winter *et al.*, 2007; Hoopes *et al.*, 2019; Woodhouse *et al.*, 2021).
